# A method for high throughput image based antifungal screening

**DOI:** 10.1101/2021.08.18.456906

**Authors:** Rosalie Sabburg, Aphrika Gregson, Andrew S. Urquhart, Elizabeth A. B. Aitken, Linda Smith, Louise F. Thatcher, Donald M. Gardiner

## Abstract

Robust antifungal screening is technically challenging particularly for filamentous fungi. We present a method for undertaking antifungal screening assays that builds upon existing broth dilution protocols and incorporates time resolved image-based assessment of fungal growth. We show that the method performs with different fungi, particularly those for which spores can be used as inoculum, and with different compound classes, can accurately assess susceptibility or otherwise in only few hours, performs well even without replication and can even account for differences in inherent growth properties of strains.

## 1 Introduction

The characterisation of antifungal activity of compounds and mixtures is challenging in both the laboratory and clinical environments. Methods largely rely on the assessment of growth after a defined, and sometimes prolonged, period of time. For example, current methods in a clinical diagnostic setting rely on naked eye visual assessment or optical density measurements of endpoint growth in microtitre plates for most fungi (Arendrp *et al*., 2017, Alastruey-Izquierdo *et al*., 2015). Although antifungal susceptibility testing is largely standardised in clinical settings, read outs can be considerably variable depending upon a number of factors such as the laboratory in which the assay is conducted, inoculum size, variation between media batches, the definition of what the end point is as well as the methodology of reading the assays (Kidd *et al*., 2021). In research laboratories spectrophotometry is typically used with endpoint or time course readings, which is a technology that has largely been adopted from bacterial growth assessment. However, problems with optical density measurements include the propensity of filamentous organisms to grow in clumps which can be exacerbated by the antifungal agents. High numbers of replicates can be used to circumvent the inherent variability, but low or no replicates would be desirable in some situation such as compound discovery work. Indeed, in the clinical setting no replicates are used to assess susceptibility/resistance to antifungals.

Image based assessment of antibiotic susceptibility in bacteria has advantages such as very rapid time frames compared to growth assessment using optical density measures (Fredborg *et al*., 2013). Other technologically advanced approaches are beginning to be reported for fungi such as reflectomic inference spectroscopy in specially fabricated devices (Heuer *et al*., 2020). In theory, image-based assessment should allow issues such as ‘non-homogeneous’ growth to be addressed without major changes (with the exception of instrumentation) to existing workflows. Here we report a method for undertaking the assessment of antifungal activity based on image analysis of fungi growing in microtitre plates in liquid medium. The method is applicable to many fungi that can be grown in axenic culture and can be used in both a drug discovery and research settings and could be applicable to clinical use with appropriate instrumentation. Along with the demonstration of the image-based method we incorporate, established and broadly applicable mathematical modelling approach (Sebaugh, 2011, Noel *et al*., 2018, Ritz *et al*., 2015) to characterising antimicrobial activity.

## 2 Methods

### 2.1 Fungal strains

*Verticillium dahliae* strain *BRIP 71171* was obtained from the Department of Agriculture and Fisheries (Queensland Australia) phytopathology herbarium. *Aspergillus fumigatus* isolate *SRRC 2006* was obtained from the CSIRO mycology collection (Sydney Australia) and is a reference strain for the International Committee on *Penicillium* and *Aspergillus*.

*Fusarium graminearum* isolate *CS3005* (Gardiner *et al*., 2014) and a derivative that carries a mutation in *Os1* known to impart resistance to fludioxonil (Gardiner & Kazan, 2018) was used. The *Os1* mutant strain was derived from a transformant that also carried a transgene cassette encoding Cas9, a gRNA targeting *Os1* and hygromycin resistance. The mutant strain was purified by back-crossing this transformant to *CS3005* and a single ascospore derived colony that was resistant to fludioxonil and susceptible to hygromycin was used for the experiments. The mutant allele carried a 276 bp deletion in *Os1*.

### 2.2 Fungal transformation

*Verticillium dahliae* was transformed with a construct for constitutive expression of enhanced green fluorescent protein (eGFP) with the transformation performed essentially as described for another fungus (Gardiner & Howlett, 2004). The construct was generated using yeast recombinatorial cloning as follows. An 833 bp fragment of the *Aspergillus nidulans* translation elongation factor (*TEF*) alpha promoter fused to *eGFP* and 274 bp of the *A. nidulans TEF* terminator synthesised as a gBLOCK by Integrated DNA Technologies (Singapore) with the 5’ extension cctcaccgcggcccatggtctagaactagtggatcc and 3’ extension ggatccgaattcgttaacaagcttgtcgacctcgag to allow recombination-based cloning in yeast. The TEF promoter-eGFP-TEF terminator fragment was cloned into the *Bam*HI site of vector pPZPnat1 (Gardiner *et al*., 2005). At the same time a 2μ origin of replication and *URA3* selectable marker were added to the backbone of pPZPnat1 (Genbank AY631958.2) in the *Sca*I site to allow selection and maintenance in yeast. The *URA3* and 2μ components were amplified from pYES2 using primers (actatagcagcggaggggttggatcaaagtcttcctttttcaatgggtaataactga and caaccacagggttcccctcgggatcaaagtacaatcttcctgtttttggggc) using Phusion DNA polymerase (New England Biolabs) with annealing at 63°C and a 2 minute extension. The resulting plasmid was extracted from yeast using the Zymoprep yeast miniprep kit version II (Zymo Research, California, USA), amplified in *E. coli* TOP10 and sequence verified before triparental mating using the helper strain RK2013 to transfer the construct into *A. tumefaciens* strain AGL1 as described elsewhere (Wise *et al*., 2006) except that the mating phase was conducted without a membrane and *A. tumefaciens* containing the construct were selected by subculturing a loop of the mating mix onto selective solid LB media containing Rifampicin (50 mg L^-1^), Ampicillin (100 mg L^-1^) and Kanamycin (50 mg L^-1^).

### 2.3 Antifungal assays

Spores for antifungal assays were prepared from each species grown on ½ PDA. Approximately 1 cm^2^ of colonised agar was excised from the plate and transferred to approximately 10 mL of ½ PDB, vortexed and filtered through a 40 μm cell strainer (Falcon®). Test compounds were dissolved in either dimethyl sulfoxide (DMSO) (tebuconazole, carbendazim, fludioxonil) or water (hygromycin, nourseothricin). Stock solutions were: tebuconazole (65 mM), carbendazim (5.23 mM), fludioxonil (100 mM), nourseothricin (83 mM) and hygromycin (95 mM). Two-fold dilutions series were set up in 0.2% DMSO (tebuconazole and fludioxonil), 2% DMSO (carbendazim) or water (hygromycin and nourseothricin). Test compound dilutions were mixed with an equal volume of spores in ½ PDB in a 96- or 384-well microtitre plate. Assays in 96 well plates were conducted in a final volume of 100 μL and in 384 well plates the final volume was 60 μL. No-treatment controls were included containing an appropriate amount of DMSO or water.

For analysis of the response of *A. fumigatus* to tebuconazole, both bright field and optical density measurements were taken after 24 hours of growth at room temperature. Automated bright field imaging was conducted using a Cytation 1 reader (Biotek) fitted with high contrast bright field imaging capability controlled by Gen5 software (Biotek). A 4× objective was used. Settings were chosen that gave sharp images in the untreated control wells. The focal height was fixed to 2700 μm corresponding to bottom of the well. Images were taken using 100 ms exposure, LED intensity of 5 and no detector gain. There was a 300 ms delay between plate movement and image acquisition. Various image process parameters were trailed and assessed for their appropriateness using the visual inspection tool in the Gen5 software but total object area measurements were relatively robust to changes in these settings. The final settings used digital phase contrast processing applied to the images using default parameters. Object area was determined using the Cellular Analysis feature of Gen5 (parameters were dark background, splitting of touching objects, object size in the range 5-100 μm, objects at the edge of the image were included and the entire image was analysed and automatic background flattening settings were used based upon 5% of the lowest pixels). Optical density measurements were made using a Perkin Elmer EnVision using an absorbance monochromator set at 595 nm, with 100 flashes used per measurement with excitation intensity set at 100% with no detector gain used.

For the analysis of *V. dahliae* growth in the presence of carbendazim, both bright field and fluorescent images were captured using a 10× objective in the same plate imager as above. Imaging settings were determined at the initiation of the culture (i.e. spores) to give sharp images. For bright field the LED intensity was 7, exposure time 153 ms with a detector gain of 1.1 using autofocus with default parameters. For fluorescence the LED intensity was 10 with a 100 ms exposure and detector gain of 2. There was a 1 second delay between plate movement and imaging. Bright field images were processed using the Cellular Analysis feature in Gen5 with splitting of touching objects and an object size range of 2 and 1×10^7^ μm with the inclusion of edge objects. Background flattening during object detection was performed using a 1 μm rolling ball diameter and image smoothing strength of 3 with the background evaluated on 5% of the lowest pixels. For fluorescent images the object size was set to be between 5 and 100 μm including edge objects with the splitting of touching objects. The appropriateness of the image processing parameters was determined by visual inspection of the counting results but changes to the parameters did not drastically alter the calculated metrics.

For the analysis of *F. graminearum* growth in the presence of fludioxonil, bright field images were captured using a 4× objective in the same plate imager as above. Imaging settings were determined at the initiation of the culture (i.e. spores) to give sharp images. The LED intensity was 5 with an exposure time of 66 ms and no detector gain. Focal height was fixed to 2757 μm. There was a 300 ms delay between plate movement and imaging. Fungal growth was quantified using the Cellular Analysis feature in Gen5 with object sum area used as a measure of total fungal material. Object sum area was calculated based on the raw bright field images with a minimum and maximum object size set to 25 and 1000 μm respectively. Touching objects were split and edge objects were included. During the analysis the background of the image was flatted using a 2 μm rolling ball diameter with a smoothing strength of 5 with the background evaluated on a background based on 5% of the lowest pixels. The appropriateness of the image processing parameters was determined by visual inspection of the counting results but changes to the parameters did not drastically alter the calculated metrics.

### 2.4 Logistic regression model of dose-response curves

The dynamics of biological inhibition are often modelled using four parameter logistic regression models which is essentially a sigmoidal function (Sebaugh, 2011) (Equation 1). This was implemented in SigmaPlot 14.5 using default parameters. Both the image and optical density measurements were modelled in the same way. Estimates of the IC_50_ (concentration at which growth is 50% of the maximum) were obtained along with the Hill Coefficient (which is the slope of the curve at the IC_50_ and describes the shape of the sigmoidal curve) and 95% confidence intervals. Data were exported to Microsoft Excel for plotting and exported to Adobe Illustrator for figure preparation. An estimate of the IC_90_ (concentration at which there is 90% inhibition of growth) was calculated in Microsoft Excel using the estimated IC_50_ and Hill coefficient using a rearranged version of the logistic function with the minimum and maximum response set to 0 and 100 respectively and response value (i.e. y) set to 10 (ie 100-90) (Equation 2).

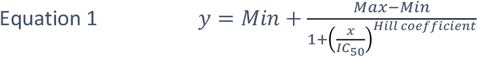

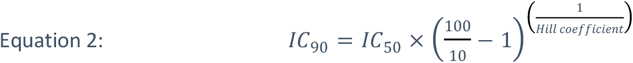

## 3 Results

### 3.1 Bright field image analysis is a suitable alternative to optical density measurements

To assess the suitability of image-based assays, a simple experiment was set up to measure the effect of the antifungal tebuconazole on the growth of *A. fumigatus* using a two-fold dilution series of the active compound from 0.032-65 μM (12 distinct concentrations). Using a single time point of 24 hours after initiation of the fungal growth both imaging and optical density measurements allowed the assessment of the inhibition by tebuconazole Figure 1. Although the calculated IC_50_ values were similar (Table 1), the shapes of the curves were different, resulting quite different IC_90_ estimates. The standard error for the predicted Hill coefficient was considerably larger for the optical density measures suggesting a greater uncertainty with this data which is also reflected in the 95% confidence intervals for the modelled curves (Figure 1B&C). Taken together, in an endpoint assay, image analysis performs at least as good as, if not better, than optical density in assessing sensitivity to an antifungal agent for the clinically important species *A. fumigatus*.

**Figure 1:**
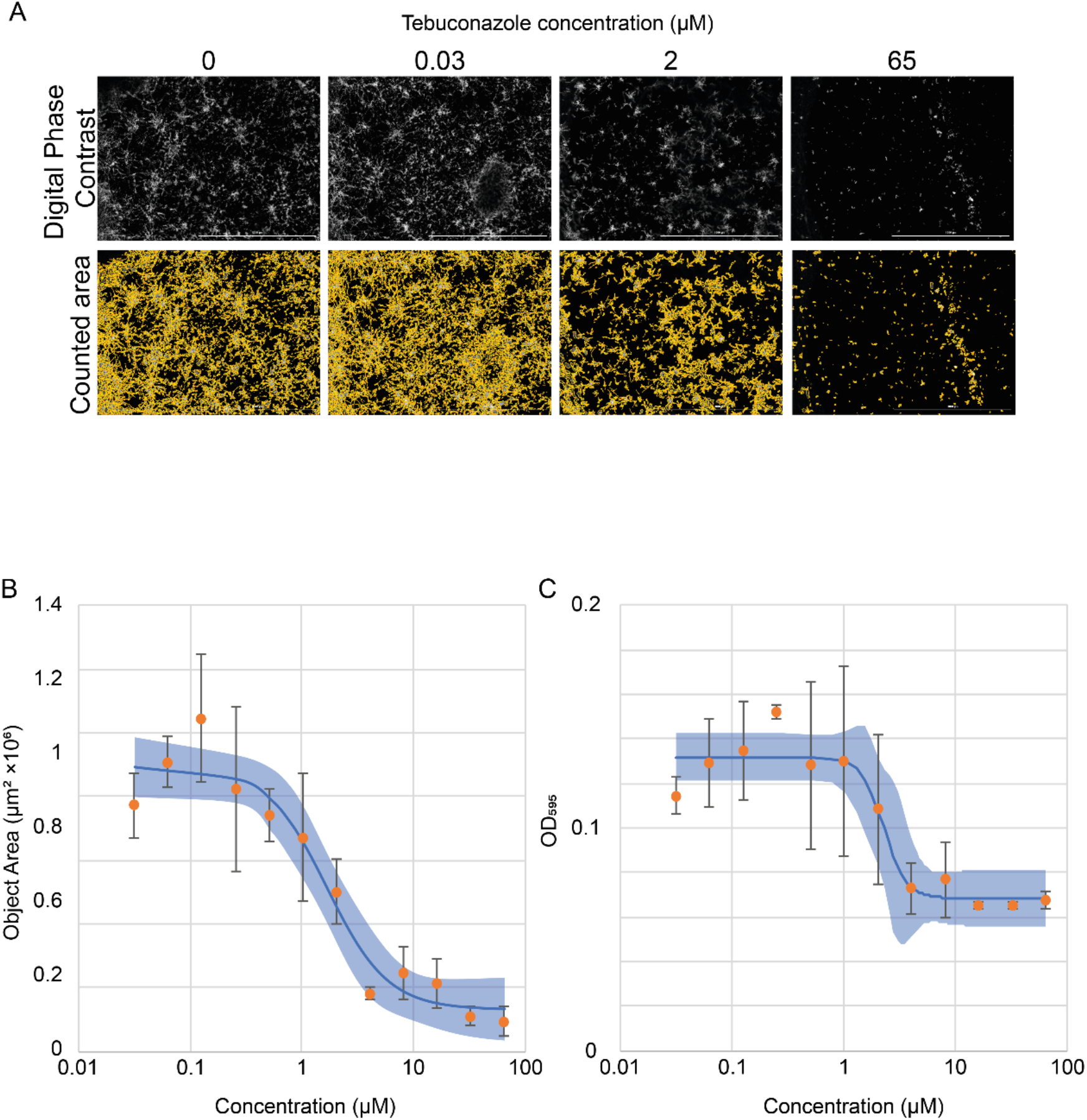
Single time point analysis (24 hours) of the effect of tebuconazole on the growth of *Aspergillus fumigatus*. (A) Exemplar digital phase contrast images of *A. fumigatus* grown in differing concentrations of tebuconazole along with the results of image analysis (yellow). (B) Image-based and (C) optical density-based dose response and modelled logistic regression curves (blue line) and 95% confidence interval (shading). Measurements are shown as the mean of three biological replicates and error bars represent the standard error. The same experimental plate was used to generate the data in the entire figure.

**Table 1:** Estimated parameters for the growth inhibition imparted by tebuconazole on *Aspergillus fumigatus* as measured by image analysis and optical density. IC50 and the Hill coefficient values are presented with the standard error.

| Parameter | Image analysis | $\text{OD}_{595}$ |
| --- | --- | --- |
| $\text{IC}_{50}$ | $1.674 \pm 0.374 \mu\text{M}$ | $2.303 \pm 0.491 \mu\text{M}$ |
| $\text{IC}_{90}$ | $6.621 \mu\text{M}$ | $3.891 \mu\text{M}$ |
| Hill Coefficient | $1.598 \pm 0.5179$ | $4.188 \pm 3.752$ |

### 3.2 Time resolved imaging allows for rapid inhibitory concentration estimation

Having established that image analysis is largely equivalent to optical density measurements for fungal growth (using *A. fumigatus* as a test organism), and in particular measurement of IC_50_, we sought to analyse in more detail the potential for earlier estimation of growth and antifungal activity. We also sought to validate the general applicability of image analysis to other fungi. To achieve this, repeated imaging was undertaken of a fluorescent *V. dahliae* strain growing in the presence of various concentrations of the fungicide carbendazim. Both bright field and fluorescence images were collected every two hours over a total of 30 hours. The fungal growth was largely confluent by approximately 16 hours in the untreated control conditions (Figure 2A, Supplementary Figure 1A and video 1) and as such the antifungal activity of carbendazim was assessed using data up to this time point. Indeed image analysis after the growth became confluent was problematic, particularly using bright field images, where the classification of image elements was less accurate and resulted in area calculations that decreased after this point (data not shown).

**Figure 2:**
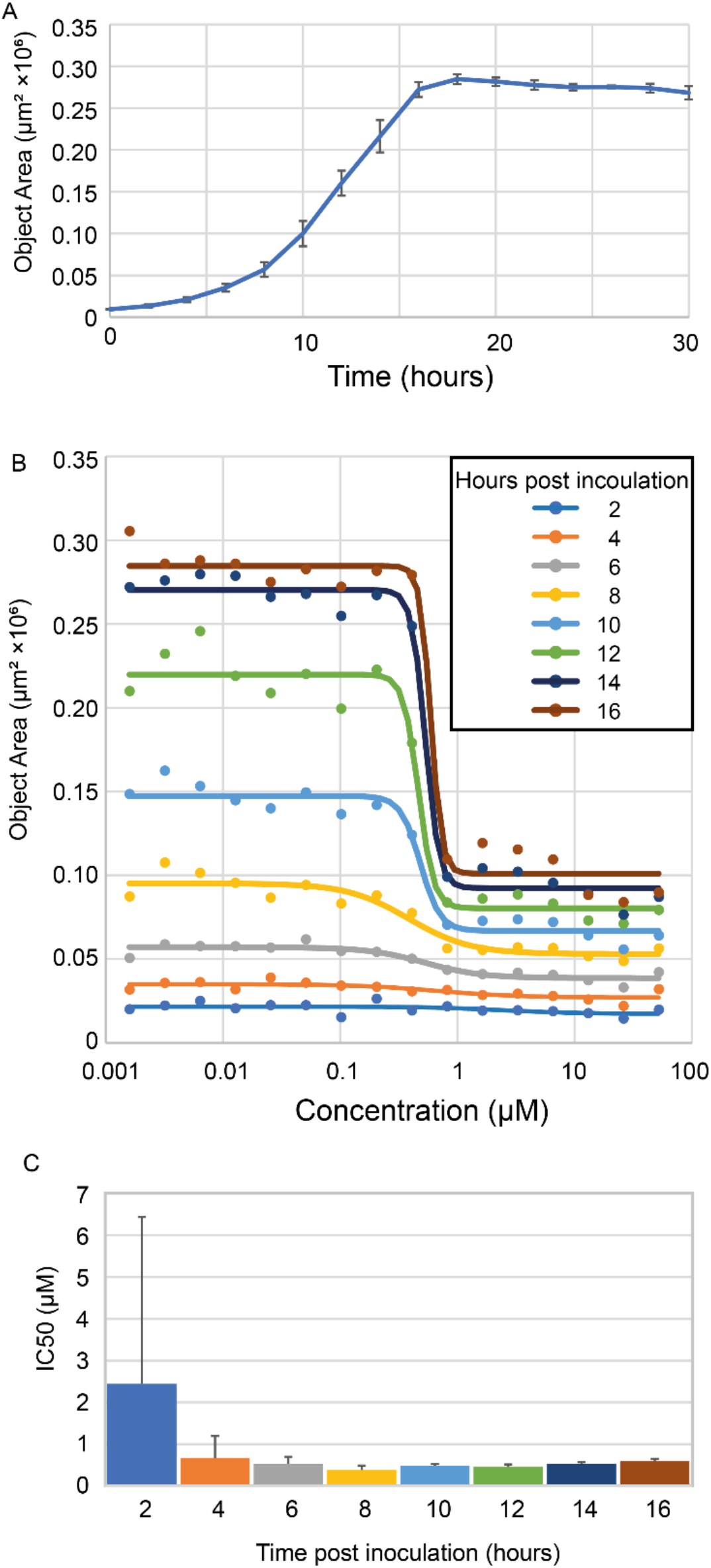
Assessment of sampling time for estimation if inhibition by the fungicide carbendazim using fluorescence data from a GFP expressing strain of Verticillium dahliae. The fungus was grown in 16 different concentrations of carbendazim plus no treatment controls and imaged every 2 hours for 30 hours. (A) GFP fluorescence data for the untreated control (0.1% DMSO) over the 30-hour sampling (B) Modelling of fungal growth inhibition by carbendazim at multiple time points after inoculation. Points are the means of measured data (four biological replicates) and the lines are the modelled logistic regression. (C) Estimated IC50 at each of the time points for which logistic regression was carried out. Errors are the standard error for the IC50 as calculated by SigmaPlot.

Although growth differences were easily discernible at the later time points (8-16 hours), the collected fluorescence data allowed estimation of IC_50_ concentration from as early as four hours post inoculation (Figure 2B&C) with the estimated IC_50_ value being in the range 0.39-0.67 from four hours onwards. Using bright field data estimates of IC_50_ were similarly possible between 6 and 12 hours post inoculation (with a range of 0.53-092) but the estimates became less reliable (although remained in the same range) using later time points (Supplementary Figure 1B&C).

**Still image of Video 1:**
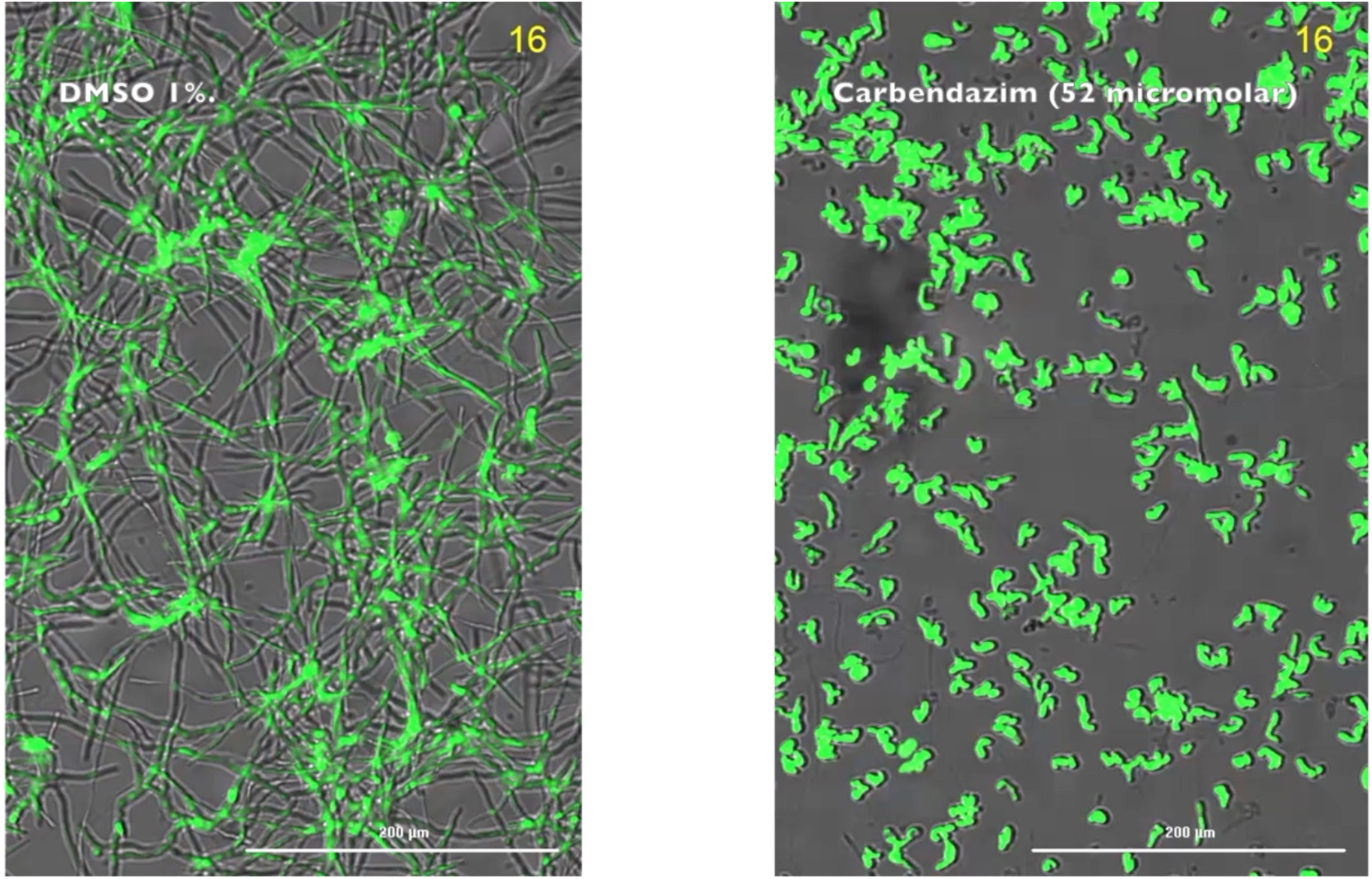
GFP expressing *Verticillium dahliae* grown for 16 hours in control 1%DMSO or 52 μM carbendazim. The bright field and GFP images were overlayed.

### 3.3 Comparative analysis of fungal strains

One of the common research applications of antifungal assays is the comparative analysis of strains or mutants. For example, comparing individuals in a field population of fungi for fungicide resistance or comparing mutant strains with their parent. These types of comparative assays can be complicated by difficulty equalising the input inoculum into the assay or by inherent differences in growth rates between strains, independent of their sensitivity to a particular antifungal agent. To assess the utility of image based analyses for comparing strains we analysed the response of a wild type *F. graminearum* and a derived mutant strain carrying a deletion of 276 bp in the osmosensor histidine kinase 1 (*FgOs1*, locus tag FG05_07118) gene to fludioxonil. Null mutations of this gene impart resistance to phenylpyrrole fungicides such as fludioxonil (Gardiner & Kazan, 2018). In this assay, despite using side by side spore preparations from identically aged source plates and use of an automated cell counter to quantify the number of spores used as inoculum, there were visibly more spores of the *Os1* strain included in assay. Moreover the *CS3005* and *Os1* germinating spores looked different (Figure 3A) and the dynamics of their growth in unamended conditions was different (Figure 3B). However, to account for this, measurements were analysed for time points at which the calculated area of fungal growth in the unamended controls was approximately half maximal and this was expressed as being relative to the growth in unamended media (Figure 3C). Although this approach should assist with comparisons between strains that vary in intrinsic growth rate, as shown above, the inhibition dynamics (i.e. IC_50_) don’t drastically differ when different time points are chosen provided some growth has occurred since the initiation of the assay. The *FgOs1* mutant strain of *F. graminearum* was highly resistant to fludioxonil (Figure 3C&D) and showed no inhibition of growth even at high concentrations of fludioxonil (12 μM), compared to the wild type for which growth was ablated above 0.4 μM. Taken together this analysis suggests that image based analysis of growth is highly suited to between strain comparisons even when their inherent growth properties differ.

**Figure 3:**
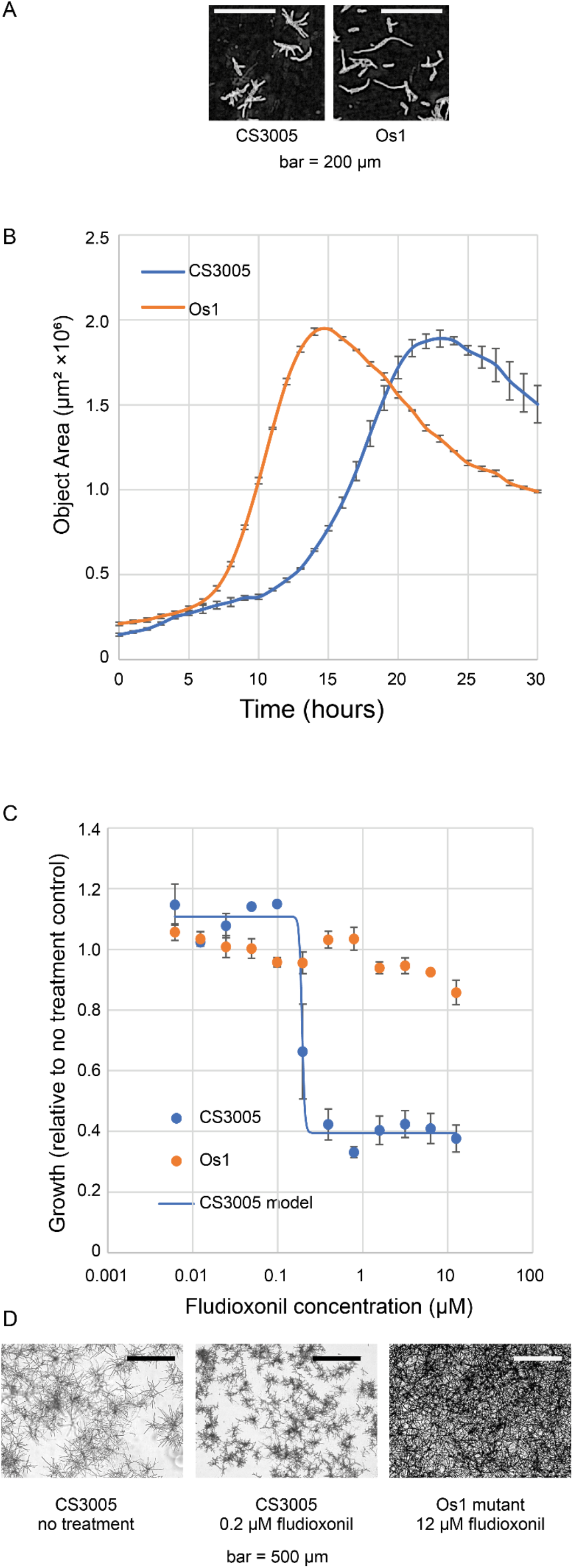
Comparison of strains for sensitivity to the antifungal fludioxonil. (A) The wild type strain (*CS3005*) and a mutant carrying a deletion in *Os1* growing in ¼ PDB containing 0.1% DMSO. The *CS3005* was taken 12 hours post inoculation and the Os1 image 8 hours post inoculation. Digital phase contrast images are shown. (B) Quantitation of growth of *CS3005* and *Os1* strains growing in ¼ PDB containing 0.1% DMSO. (C) Strains were grown in a range of fludioxonil concentrations and growth expressed as being relative to an unamended control treatment. Error bars represent the standard error of the mean of three biological replicates. Data represented are from imaging at 17 and 10 hours post inoculation for *CS3005* and *Os1* respectively. *CS3005* showed severe sensitivity to fludioxonil at concentrations above 0.2 μM. (D) Exemplar bright field photographs of the parental strain (*CS3005*) growing without fludioxonil, *CS3005* growing in fludioxonil at the approximate IC_50_ and the *Os1* mutant growing at a high concentration of fludioxonil. Images are from 20 hours post inoculation.

### 3.4 Replication is not essential

The need for multiple serial dilutions in measuring the growth inhibition properties of antifungal agents means there is a limited capacity using standard format microtitre plates to compare large numbers of strains or compounds in parallel. We sought to experimentally address how important replication is in these types of experiments. Indeed, four parameter logistic regression with an unreplicated dilution series can still be used to estimate the error associated with each of the parameters. To test if this was appropriate for biological samples such as these, we analysed replicated dilution series in assays assessing the antifungal properties of three different test compounds (hygromycin, nourseothricin and carbendazim) as unreplicated datasets and compared the generated models to that obtained when the replicates were combined into a single regression. As shown in Figure 4, for all compounds the estimation of IC_50_ showed minimal differences whether replication was used or not. However, the shapes of the regression models differ somewhat and this was particularly pronounced for one individual dilution series in the carbendazim datasets.

**Figure 4:**
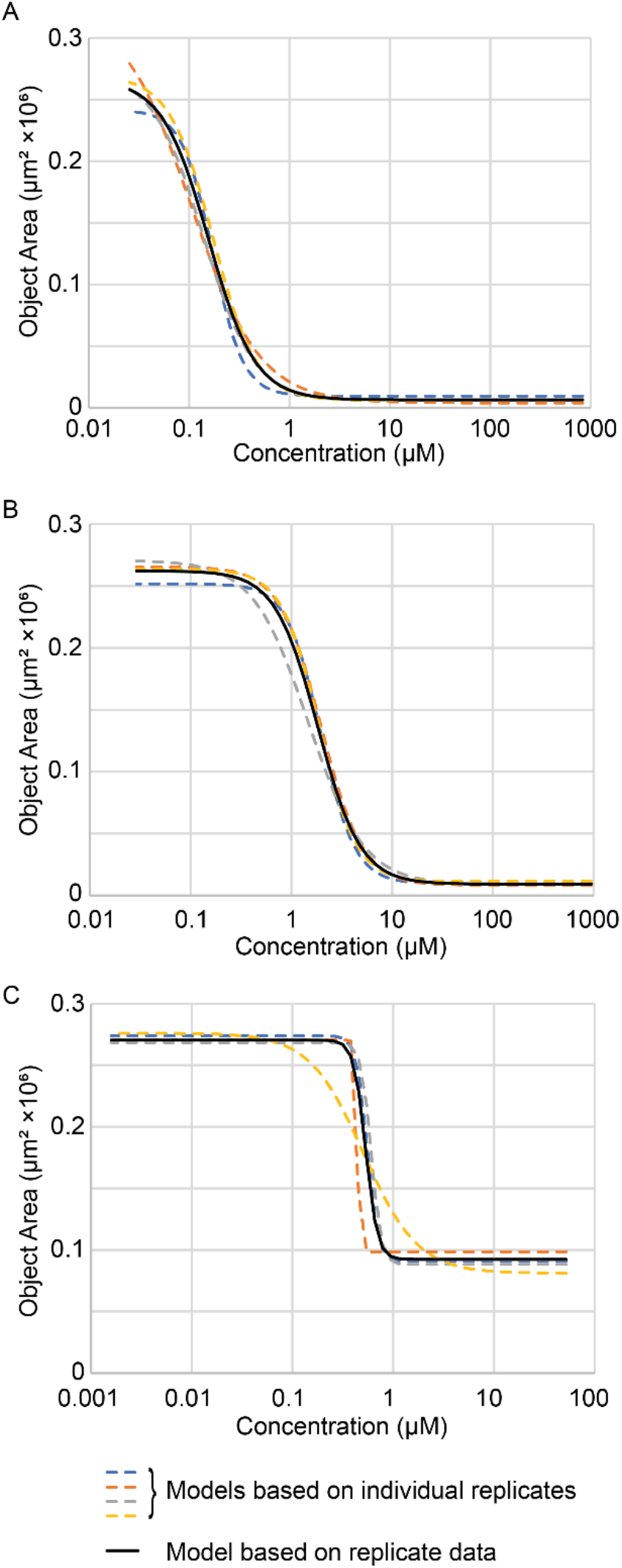
Analysis of the importance of replication in determining inhibition of fungal growth using serial dilution of test compounds and image based quantification of fungal growth. Three different compounds were tested; hygromycin (A), nourseothricin (B) and carbendazim (C). Four replicate dilutions series were assessed both individually and in a combined model.

Analysis of single dilution series from the tebuconazole inhibition of *A. fumigatus* data presented above (Figure 1), with both bright field imaging and optical density measurements, were also undertaken. Similar estimates of IC_50_ were obtained with the bright field image data from all three replicates (Figure 5A). However for optical density measurements from the same biological material analysis of one individual dilution series resulted in an IC_50_ estimation that was ∼5-fold lower than that estimated by the replicated data (Figure 5B). Given the variability associated with the OD values (as presented for this data in Figure 1C) this is not surprising. Taken together replication of dilutions series is not absolutely critical, except where finer estimates of IC_50_ and in particular IC_90_ are required. Image analysis may perform slightly better than optical density measurements in the absence of replication.

**Figure 5:**
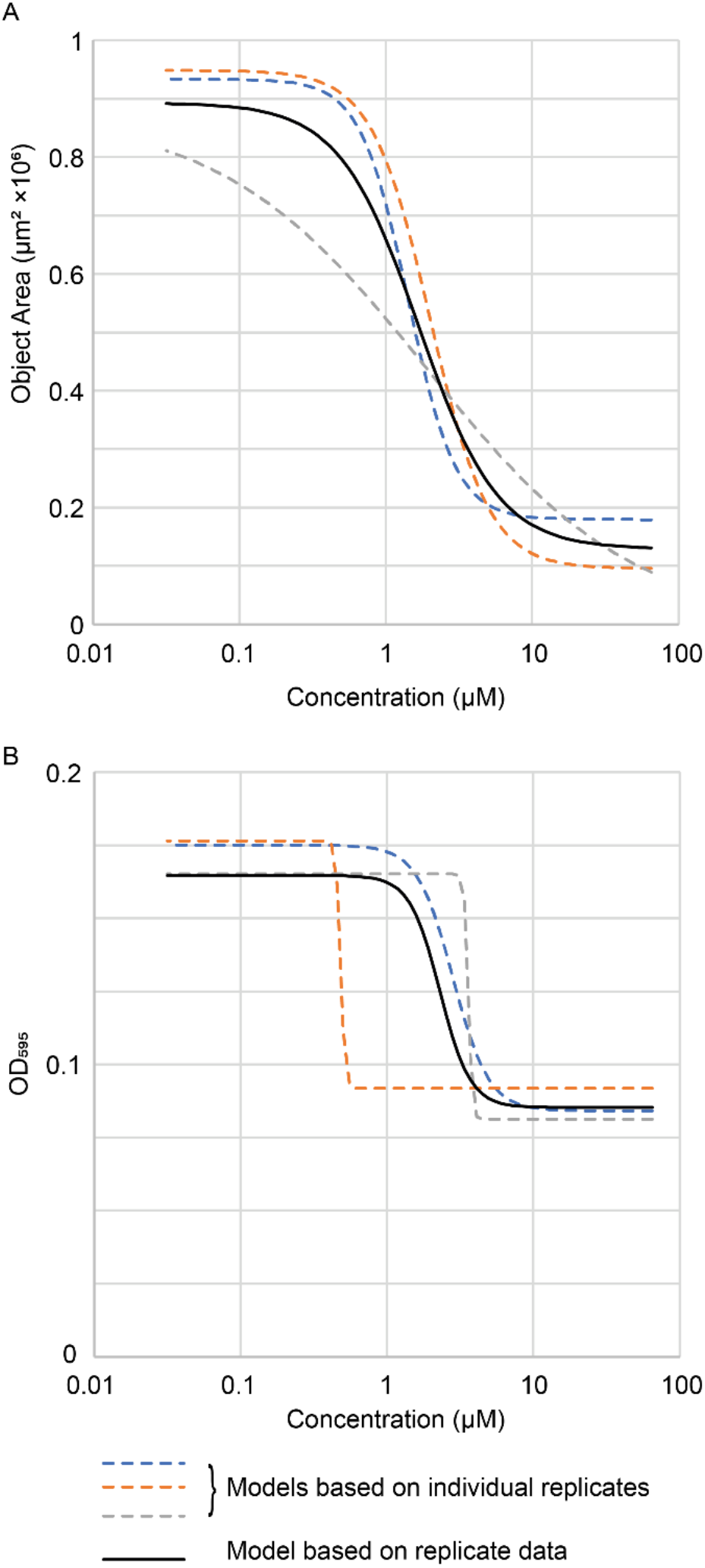
Analysis of the importance of replication in determining inhibition of fungal growth using serial dilution of test compounds and (A) image- or (B) optical density-based quantification of fungal growth. The data presented in Figure 1 for *Aspergillus fumigatus* grown in the presence of tebuconazole were used here.

## 4 Discussion

Estimation of antifungal activity is technically challenging. Adding to the technical challenges, the commonly utilised measure of growth (optical density or even visual assessment) (Arendrp et al., 2017, Alastruey-Izquierdo et al., 2015) has limitations that we here show can be somewhat circumvented with image-based analysis of growth. Image analysis is particularly useful for reducing the time required to estimate growth inhibition as well as providing an ability to compare between strains with vastly different growth dynamics. Analyses of growth inhibition are also aided considerably by having multiple time points to analyse. In theory time-resolved analysis should be possible with the appropriate instrumentation for both image-based and optical density-based analyses but image-based analysis is far more sensitive than optical density measures which typically is difficult without at least overnight incubation.

Data analysis is also not intuitive for all operators and indeed may require purchase of proprietary software unless the user is proficient in R (Ritz et al., 2015). Whilst we utilised SigmaPlot to implement four parameter logistic regression, equivalent analysis can be performed using a freely available online tool (AAT Bioquest, 2021). Although for many purposes four parameter logistic regression is likely to suffice, it should be noted that for some fungicides hormetic effects (growth promotion at low doses) will require a different regression model (Noel et al., 2018) but similar experimental setups should suffice.

Despite the improvements that image analysis can provide, this may not be suitable for all organisms. In our hands the most challenging component of image-based analysis of growth was ensuring the images were taken in focus. Of the three organisms tested, *A. fumigatus, F. graminearum* and *V. dahliae*, this was found to be to be easiest with *V. dahliae* which tends to grow predominantly, at least initially, on the bottom of the microtitre plates. *A. fumigatus* spores appeared to grow as a mixture of both unattached floating clumps and on the bottom of the well but with appropriate imaging parameters the growth on the well bottom could be assess with minimal interference from the unattached clumps on the surface. As time progresses *F. graminearum* creates a three-dimensional hyphal network in liquid culture that extends above the liquid-air interface, which again made imaging difficult at the later time points. We also tested our imaging methodology with *Sclerotinia sclerotiorum* using hyphal fragments as input material, due to the absence of vegetative spores produced by this organism (Bolton *et al*., 2006). However, we were unable to reliably autofocus on this organism in culture using bright-field or even when using a GFP expressing strain (data not shown) and as such couldn’t quantify the growth with any certainty.

In summary, we present a robust yet flexible approach to the assessment of antifungal activity that should be applicable to many (but not all) fungi, particularly in laboratory settings, for applications such as compound discovery or fungicide resistance assessment. With appropriate instrumentation the method is also amenable to automation at large scale.

## 5 Acknowledgements

Aphrika Gregson was supported by a honours scholarship awarded by the Cotton Research and Development Corporation.

## 6 Supplementary Figures

**Supplementary Figure 1:**
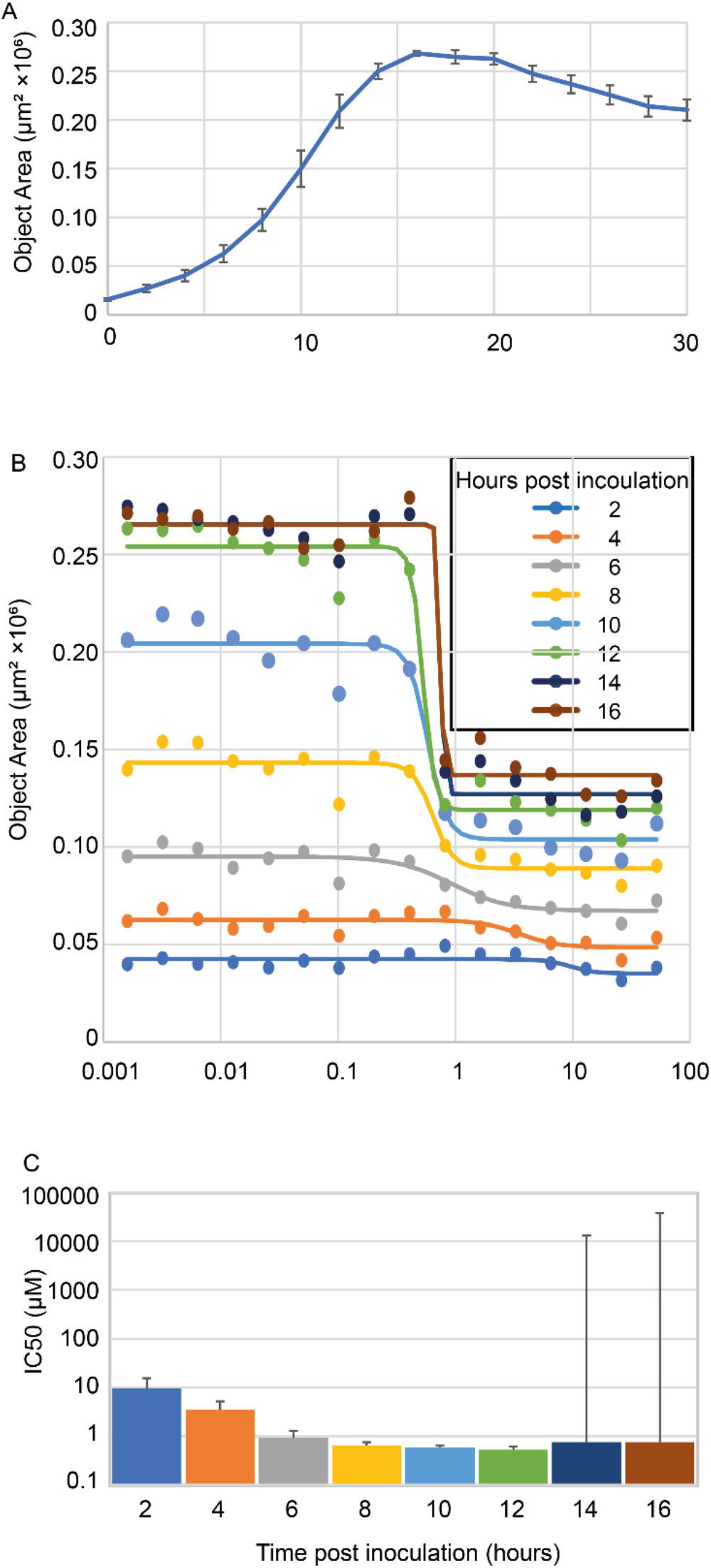
Assessment of sampling time for estimation if inhibition by the fungicide carbendazim using brightfield image data from a GFP expressing strain of Verticillium dahliae. The fungus was grown in 16 different concentrations of carbendazim plus no treatment controls and imaged every 2 hours for 30 hours. (A) GFP fluorescence data for the untreated control (0.1% DMSO) over the 30 hour sampling (B) Modelling of fungal growth inhibition by carbendazim at multiple time points after inoculation. Points are the means of measured data (four biological replicates) and the lines are the modelled logistic regression. (C) Estimated IC50 at each of the time points for which logistic regression was carried out. Errors are the standard error for the IC50 as calculated by SigmaPlot.

